# biscot: an Optimal Transport framework for multimodal bacterial single-cell data analysis

**DOI:** 10.1101/2025.03.28.645895

**Authors:** Medina Feldl, Gorkhmaz Abbaszade, Florian Schattenberg, Kathrin Stückrath, Susann Müller, Christian L. Müller

**Author notes:** **Correspondence** All correspondence and requests for material should be addressed to C.L.M.

## Abstract

Computational optimal transport-based approaches have emerged as promising tools for the integration and interpretation of complex single-cell data. In this study, we introduce an integrative Optimal Transport (OT) framework for spatiotemporal and multi-omics *bacterial* single-cell analysis using Gaussian Mixture Model (GMM) OT, termed biscot (bacterial integrative single-cell optimal transport). We show that biscot, equipped with a novel global-to-local GMM initialization, outperforms classical OT and entropically-regularized OT methods both in terms of speed and accuracy for disentangling complex bacterial communities mixtures from single-cell flow cytometry data. When applied to time-series flow cytometry data from *Bacillus subtilis*, our framework delivers robust and biologically meaningful results, effectively capturing subtle phenotypic shifts in spore populations transitioning from inactive to active growth states. `biscot` also allows multi-omics integration of flow cytometry and unpaired bacterial single-cell RNA sequencing (scRNA-seq) data, enabling the alignment of individual gene expression profiles to the cytometric data. For an unpaired flow cytometry/scRNA-seq dataset of *Bacillus subtilis* cells, we validate the biological plausibility of inferred gene expression patterns with relevant marker genes, including *spoVID* and *nin*, closely aligning with observed cellular states. Overall, our framework thus provides not only dynamic tracking of phenotypic cell states but aligns cell states with detailed transcriptomic information from scRNA-seq, demonstrating its potential to advance microbial single-cell research. `biscot` will be made publicly available on GitHub.

## Introduction

Flow cytometry is an essential tool for single-cell analysis in microbiology, enabling high-throughput characterization of bacterial heterogeneity at single-cell resolution [1]. By capturing multivariate phenotypic signals, including cell properties derived from light scattering and dye-dependent intracellular components, it allows simultaneous profiling of thousands of cells, facilitating the study of complex microbial communities and their adaptive responses to environmental changes. However, traditional methods of flow cytometry analysis, such as manual gating and histogram-based comparisons, often do not fully capture the rich multi-dimensional structures in cytometry datasets and, when time-series or multi-condition data are available, provide only coarse-grained snapshots that limit our ability to compare and track dynamic bacterial subpopulations over time or environmental gradients.

Computational techniques based on Gaussian Mixture Models (GMMs) have been developed to improve subpopulation identification in bacterial flow cytometry. Algorithms such as `flowClust` [2], `flowPeaks` [3], `flowEMMI` [4], and `PhenoGMM` [5] automate the gating of cell populations, refine phenotypic characterization, and improve reproducibility. While these methods offer improved clustering, they typically operate on individual cytograms (e.g., measured at single time points or at specific conditions), followed by aligning and merging individual gate templates or clusters via algorithmic matching [6]. Furthermore, the majority of tools and workflows are dedicated to data from flow cytometric measurements, limiting the integration of other emerging single-cell technologies.

While flow cytometry provides cost-effective high-throughput single-cell phenotypic information, recent advances in bacterial single-cell RNA sequencing (scRNA-seq) offer complementary insights through high-resolution transcriptomic profiling, albeit at higher cost and lower throughput. The integration of these two modalities could, in principle, provide deeper insights into microbial heterogeneity and dynamics, yet faces both conceptual and computational challenges due to the *unpaired* nature of the multi-omics measurements (i.e., each technology operates on a disjoint set of cells), differences in measurement scales, sparsity levels of the underlying primary data, and distinct (sub-)population distributions.

Optimal Transport (OT) frameworks offer promising solutions to such integration challenges and have been heavily adopted in eukaryotic single-cell analysis [7, 8]. The principle idea behind OT methods is to conceptualize single-cell measurements as multi-dimensional distributions over a (potentially latent) space and cast the problem of comparing and tracking cells as distributional transport problem where one distribution is warped into another one at minimum cost. Key approaches include (i) classical Wasserstein or Earth Mover’s Distance (EMD) OT for direct cell-level alignment [9, 10], (ii) entropically regularized OT [11], (iii) Gromov-Wasserstein OT approaches [12], and (iv) Gaussian Mixture Model-based OT (GMM-OT) [13, 14]. These and related approaches have found numerous applications in human single cell research [15]. For instance, `Waddington-OT` [7] uses EMD to estimate cell differentiation trajectories from clustered single-cell data. Similarly, `scEGOT` [16] introduced an OT framework using GMMs with entropic regularization for trajectory inference in eukaryotic scRNA-seq data. `QOT` [17] applied GMM-OT to compute distance matrices for large RNA-seq datasets, dynamically capturing gene expression patterns across samples with improved computational efficiency. The `scot` framework [12] uses Gromov-Wasserstein OT to align different unpaired single-cell modalities. Finally, the `moscot` framework [8] allows both trajectory inference and multi-omics data integration and provides multiple computational improvements, including near-linear run time complexity through entropic regularization, GPU acceleration, and online cost function evaluation. Notably, OT approaches to flow cytometric data have been far less pervasive. The `optimalFlow` framework [18] uses Wasserstein distances for cytometric gating and matching. Similarly, `CytOpt` [19] estimates cell population proportions from flow cytometry data using OT in a supervised fashion that can include side information (domain adaptation) in the process. However, existing OT methods are predominantly tailored to eukaryotic datasets and do not address the unique challenges of bacterial single-cell data: extreme population imbalances, high technical variability, rapid temporal shifts, and pronounced growth phase transitions. These bacterial-specific challenges require specialized computational approaches that can handle unpaired multimodal integration while maintaining biological interpretability.

In this work, we present `biscot` (bacterial integrative single-cell optimal transport), a computational GMM-based Optimal Transport framework for bacterial single-cell data analysis and integration. `biscot` includes several features tailored toward single-cell flow cytometry data, including a global-to-local initialization strategy for per-sample GMM estimation that facilitates robust tracking of bacterial population shifts via optimal transport. Furthermore, our framework enables the integration of other unpaired single-cell modalities, most prominently scRNA-seq data. We systematically evaluated our method against several OT-based approaches using the CHIC dataset [20], which provides reliable ground truth cytometric fingerprinting of complex bacterial mixtures. When applied to time-series flow cytometry datasets from *Bacillus subtilis*, `biscot` successfully tracked subtle but biologically meaningful phenotypic shifts within bacterial populations across growth phases. Furthermore, by integrating these cytometry measurements with bacterial scRNA-seq data obtained at corresponding optical densities, `b-scot`‘s transport maps enable the assignemnt of gene expression profiles to individual flow cytometry cells, thus allowing the characterization of transcriptomic profiles associated with bacterial differentiation.

By enabling precise mixture discrimination, robust subpopulation tracking, and accurate gene expression imputation, this approach opens new avenues for studying microbial adaptation and community dynamics at unprecedented resolution, with potential applications in bacterial multicellularity, biofilm formation, and antibiotic resistance studies. `biscot` will be made publicly available on GitHub.

## Methods

We first summarize the mathematical and computational components of `biscot` and relevant background on alternative OT methods. We then detail two workflows that allow (i) bacterial subpopulation tracking across multiple (time-series) samples and (ii) multimodal data integration via optimal transport.

### Gaussian Mixture Modeling (GMM)

We employ Gaussian Mixture Models (GMMs) to represent the complex, multimodal distributions that arise from single-cell technologies, including (bacterial) flow cytometry [2] and (low-dimensional embeddings of) single-cell RNA sequencing data [21]. Formally, a Gaussian Mixture Model represents data as a weighted sum of *k* Gaussian distributions, each defined by a scalar weight *w*_*i*_, mean vector *µ*_*i*_ ∈ ℝ^*d*^, and covariance matrix Σ_*i*_ ∈ ℝ^*d*×*d*^:

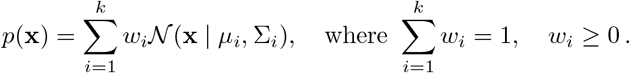

The number of components *k* is a hyperparameter that can be chosen in a data-driven way via model selection schemes or set by the expert user.

#### Parameter Estimation via Expectation-Maximization (EM)

We employed the Expectation-Maximization algorithm [22] to iteratively optimize the parameters of the GMM. Given a single-cell data matrix *X* ∈ ℝ^*n*×*d*^, the EM algorithm maximizes the log-likelihood function defined as:

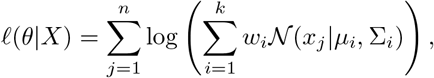

where *x*_*j*_ represents individual data points. The iteration proceeded until convergence, defined as a relative improvement in log-likelihood below a threshold of 10^−3^, or until a maximum of 1000 iterations was reached. The GMM estimation utilized the package `sklearn.mixture` (v1.6.1)[23] for all numerical computations.

#### Global-to-Local initialization Strategy

Fitting GMMs to data via maximum log-likelihood estimation is a non-convex optimization problem whose solution depends on the initialization of the optimization problem, i.e., the initial values of the component parameters. A common setting in single cell analysis is the presence of multiple samples, e.g., measurements across time or under different conditions. Our global-to-local initialization strategy consists of pooling all available samples and performing a global GMM fit on the pooled samples. Subsequently, per-sample local GMMs are fit with initial parameters derived from a global GMM. This global-to-local initialization strategy promotes consistent component labeling across multiple datasets, improving interpretability and biological relevance of the resulting clusters. We detail this approach in the Section **Multi-sample comparison of bacterial populations with** `biscot` below.

### Optimal Transport (OT)

#### Classical Optimal Transport (Earth Mover’s Distance)

Given two discrete probability measures, *µ*_*s*_ (source) and *µ*_*t*_ (target), supported on discrete point sets 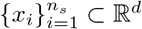 and 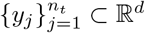, respectively:

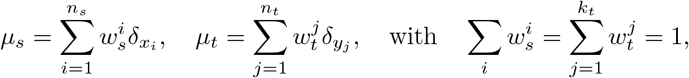

we define the OT problem as a linear program minimizing the transportation cost given by a cost function *c*(*x*_*i*_, *y*_*j*_), typically chosen as the squared Euclidean distance. The cost matrix *M* is computed element-wise based on the squared Euclidean distance:

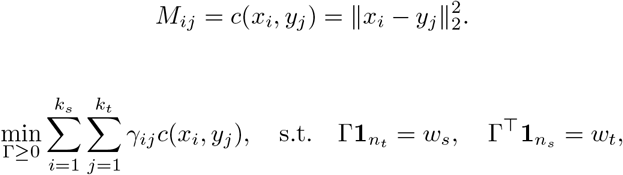

with solution Γ^*^ referred to as the optimal transport plan, and the optimal cost as the Earth Mover’s Distance (EMD)[10, 11].

### Entropic Regularization (Sinkhorn Algorithm)

To enhance the scalability of Optimal Transport (OT) computations for large single-cell datasets, we applied entropic regularization. Given two discrete probability distributions, *µ*_*s*_ (source) and *µ*_*t*_ (target), the Sinkhorn-regularized OT problem is defined as:

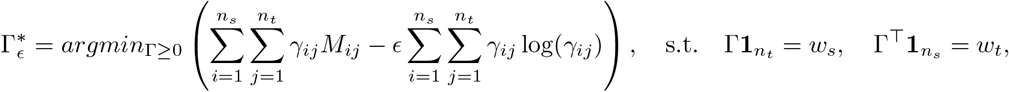

To efficiently solve this optimization, the Sinkhorn–Knopp algorithm introduces a kernel matrix *K*, defined element-wise as:

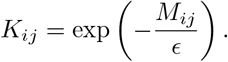

The optimal transport plan 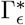 admits a diagonal-scaling factorization:

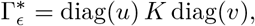

where the scaling vectors 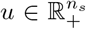 and 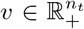 are iteratively updated to satisfy the marginal constraints. Specifically, starting with 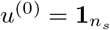 and 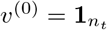, the algorithm proceeds as follows:

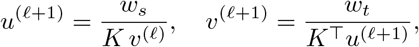

with all divisions performed element-wise. Iterations continue until convergence, typically determined when the change in *u* and *v* between successive iterations falls below a numerical tolerance (here set to 10^−9^).

Upon convergence, the resulting optimal transport plan is explicitly given by:

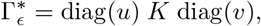

and the associated transport cost—known as the Sinkhorn distance—is computed as:

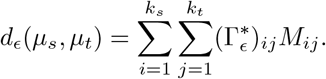

We chose the regularization parameter *ϵ* = 0.1 empirically, striking a careful balance between computational efficiency, numerical stability, and fidelity to the classical OT solution [11].

### Gaussian Mixture Model-based Optimal Transport (GMM-OT)

To extend Optimal Transport to distributions represented by Gaussian Mixture Models (GMM), we define a suitable OT framework that aligns mixtures of Gaussian distributions [14]. Consider two GMMs, the source distribution *µ*_*s*_ and the target distribution *µ*_*t*_, defined as follows:

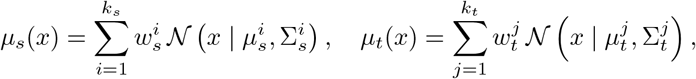

where each Gaussian component is characterized by a weight *w*, mean vector *µ*, and covariance matrix Σ.

To quantify the transport cost between individual Gaussian components from the source and target, we employ the squared 2-Wasserstein distance between Gaussian distributions:

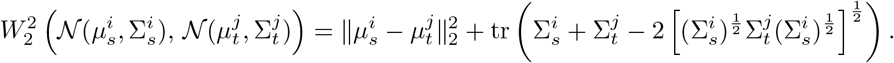

This defines the cost matrix 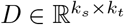, with elements:

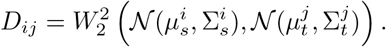

The GMM-OT problem then becomes a linear optimization that identifies an optimal transport plan 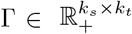, minimizing the total transport cost between Gaussian components:

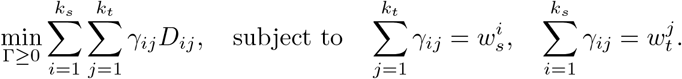

#### Barycentric Projection (Component-wise Mapping)

To translate the OT solution into explicit mappings between source and target data points, we employ the barycentric projection. For each pair of Gaussian components (*i, j*), we define an affine transformation:

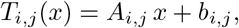

where the linear transformation *A*_*i,j*_ and translation vector *b*_*i,j*_ are calculated explicitly by:

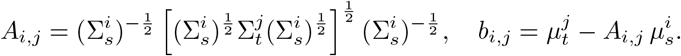

Finally, the barycentric projection, mapping any point *x* from the source distribution onto the target, is computed as the weighted sum of affine transformations:

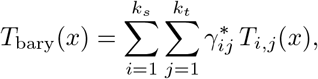

with 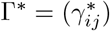 being the optimal transport plan obtained from solving the GMM-OT linear program. This projection provides biologically interpretable cell correspondences, enabling detailed exploration of cellular transitions across datasets.

### Multi-sample comparison of bacterial populations with biscot

To capture dynamic changes in bacterial populations, `biscot` combines Gaussian Mixture Models (GMMs) with global-to-local initialization and Optimal Transport (OT). We exemplify our first workflow using single cell flow cytometry data collected at multiple time points or conditions. Firstly, all data are preprocessed to remove debris, background noise, and artifacts that could interfere with downstream analyses (Figure 1, Panel 1). The cleaned datasets are then combined into a single global dataset to which a GMM is fit with a fixed number of components (Figure 1, Panel 2). In our downstream application, we choose *k* = 15, determined through benchmarking experiments. This global GMM provides a unified set of component parameters (means, covariances, and mixture proportions) that serve as a common reference frame across all time points or conditions.

**Figure 1:**
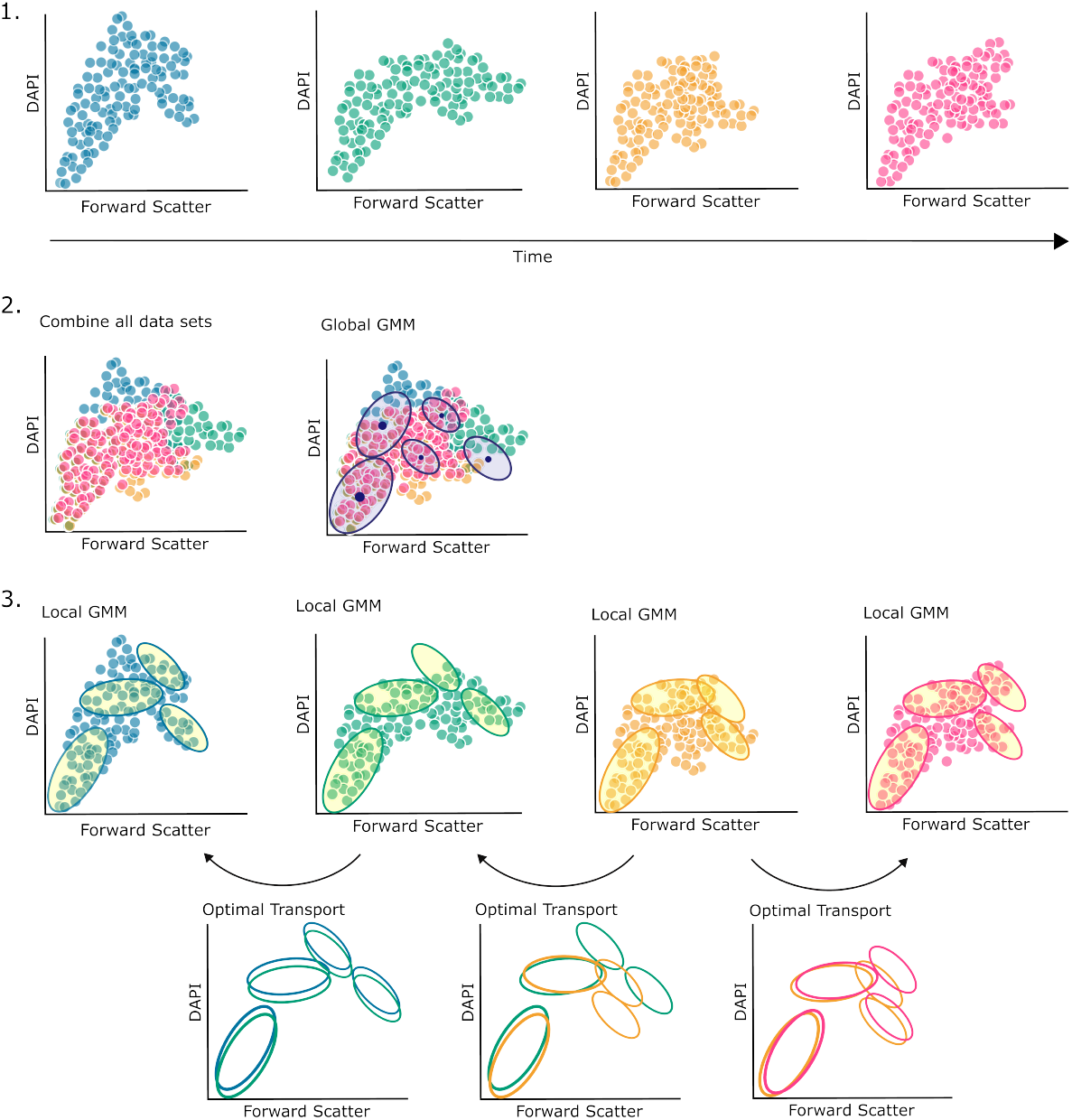
Illustration of the Gaussian Mixture Model Optimal Transport (GMM-OT) workflow applied to bacterial flow cytometry data collected at multiple time points. Initially, datasets from all time points are pooled to fit a global GMM, which ensures stable initialization across datasets. Subsequently, local GMMs are independently fitted to each dataset, preserving global consistency. Pairwise Wasserstein Optimal Transport is then incrementally applied, either forward or backward in time, anchored by manually gated reference datasets. This approach generates continuous trajectories of cell populations, facilitating the detailed analysis of phenotypic shifts within bacterial communities.

Next, we initialize local GMMs for each time point using these global parameters and then refine them by fitting to their respective individual data (Figure 1, Panel 3). This global-to-local strategy ensures that each local model inherits the same component labels while adapting to time-specific variations, enabling meaningful cross-time comparisons of bacterial subpopulations through consistent parameterization.

For time-series datasets where expert biological knowledge is available, we incorporate manual gating information from a reference time point as an anchor for the Optimal Transport procedure. Using this reference point, `biscot` computes the pairwise Wasserstein distances between consecutive time points to guide incremental OT mappings either forward or backward in time. By applying these mappings sequentially, we derive a continuous transport trajectory for each bacterial subpopulation, allowing precise tracking of phenotypic shifts that may be subtle but biologically significant (Figure 1). This approach provides advantages over snapshot-based analyses by explicitly modeling the continuous nature of bacterial population dynamics over time (Figure 1, Panel 3).

### Multimodal single-cell data integration with biscot

`biscot`‘s use of GMMs enables a seamless way to integrate multimodal data as long as individual single cell data modalities can be clustered via GMMs of the same dimension. Here, we exemplify a workflow integrating scRNA-seq and flow cytometry data which addresses fundamental differences between these modalities, including dimensionality (thousands of genes vs. few cytometric parameters), measurement scale, and data sparsity patterns (see Figure 2 for illustration of the workflow.)

**Figure 2:**
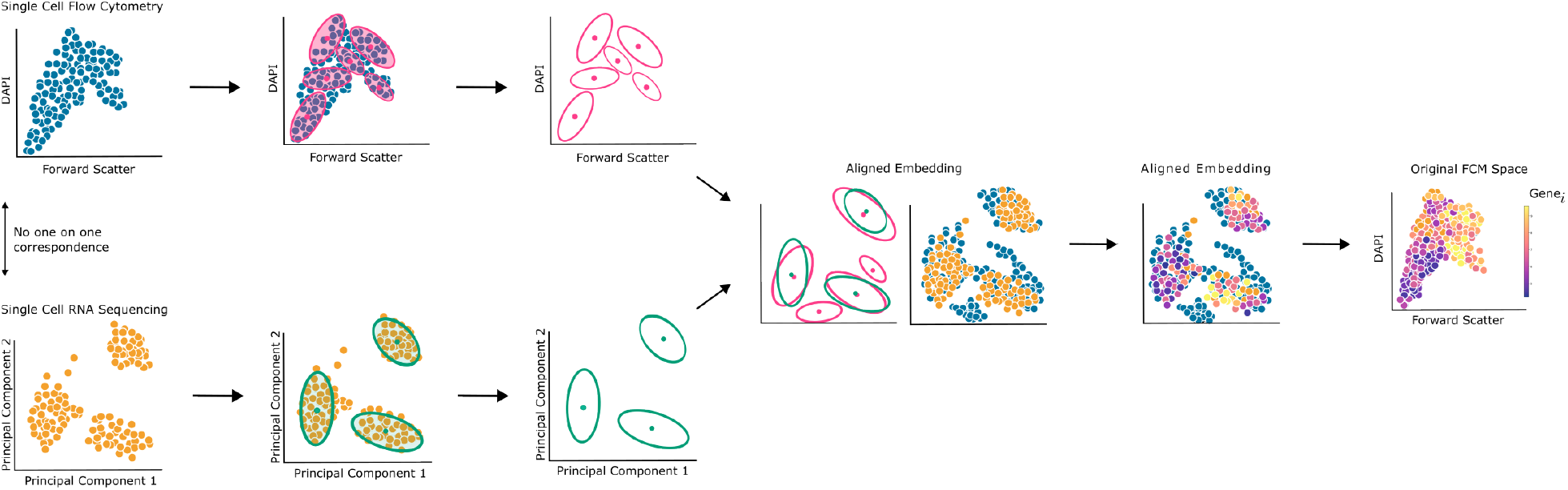
Schematic representation of the gene expression imputation workflow integrating single-cell RNA sequencing (scRNA-seq) with flow cytometry data via Gaussian Mixture Model-based Optimal Transport (GMM-OT). Following modality-specific pre-processing, separate GMMs are independently fitted to scRNA-seq and flow cytometry datasets. Pairwise Wasserstein distances between GMM components align these datasets into a shared embedding space. Gene expression profiles are subsequently imputed onto flow cytometry cells by nearest-neighbor mapping within this aligned space, thereby enriching phenotypic data with transcriptomic information.

For flow cytometry data, we applied standard pre-processing procedures including log transformation and removal of debris and mock communities. These steps harmonized measurement scales and improved the interpretability of phenotypic signals [24]. For scRNA-seq data, we employed the bacterial-specific `BacSC` pipeline [25], which addresses challenges unique to bacterial single-cell transcriptomics data: extreme sparsity, limited sequencing depth, and high technical variability. `BacSC` systematically stabilizes variance, corrects for sparsity biases, and ensures consistent scaling across genes. Following pre-processing, `BacSC` performs principled dimensionality reduction on bacterial scRNA-seq data using singular value decomposition (SVD) combined with count splitting (data thinning), learning the optimal number of latent dimensions in a data-driven fashion. Notably, for bacterial scRNA-seq data, learned optimal dimensionalities range between one and four [25].

`biscot` then independently fits local Gaussian Mixture Models (GMMs) to both single-cell datasets, effectively representing heterogeneous subpopulations within each modality. Between these fitted GMMs, `biscot` computes pairwise Wasserstein distances for corresponding components and solves the resulting GMM-OT optimization problem to generate an optimal alignment. This alignment creates a common embedding space that positions each flow cytometry cell relative to the distribution of single cells comprising gene expression profiles.

Within this aligned embedding space, `biscot` assigns gene expression profiles to each flow cytometry cell using k-nearest neighbor imputation (k=1), selecting the transcriptomic profile from the closest scRNA-seq cell in the embedding. Finally, we projected these imputed gene expression profiles back into the original flow cytometry measurement space, enriching the original phenotypic measurements with inferred transcriptomic information. This reverse projection preserves the interpretability of results within the familiar cytometric context while adding the depth of gene expression information (Figure 2). The success of this approach depends on the biological assumption that cells with similar phenotypic properties (as measured by flow cytometry) are likely to share similar transcriptomic states.

## Results

### Benchmarking Optimal Transport Methods with Cytometric Fingerprinting Data

To rigorously evaluate optimal transport (OT) methods for bacterial flow cytometry data, we used the established benchmark dataset of Koch et al. [20]. This dataset consists of three pure bacterial cultures (A, B, C) and pairwise mixtures at ten different ratios (100%, 99.5%, 95%, 85%, 70%, 50%, 30%, 15%, 5%, and 0.5%), each replicated three times, totaling 90 datasets (Figure 3). After noise and debris removal, each dataset contained on average 193,656 cells. The dataset’s uniform value ranges across samples required no additional preprocessing. Our primary benchmarking objectives were to (1) reliably identify biological replicates and (2) accurately discriminate subtle differences in mixture ratios.

**Figure 3:**
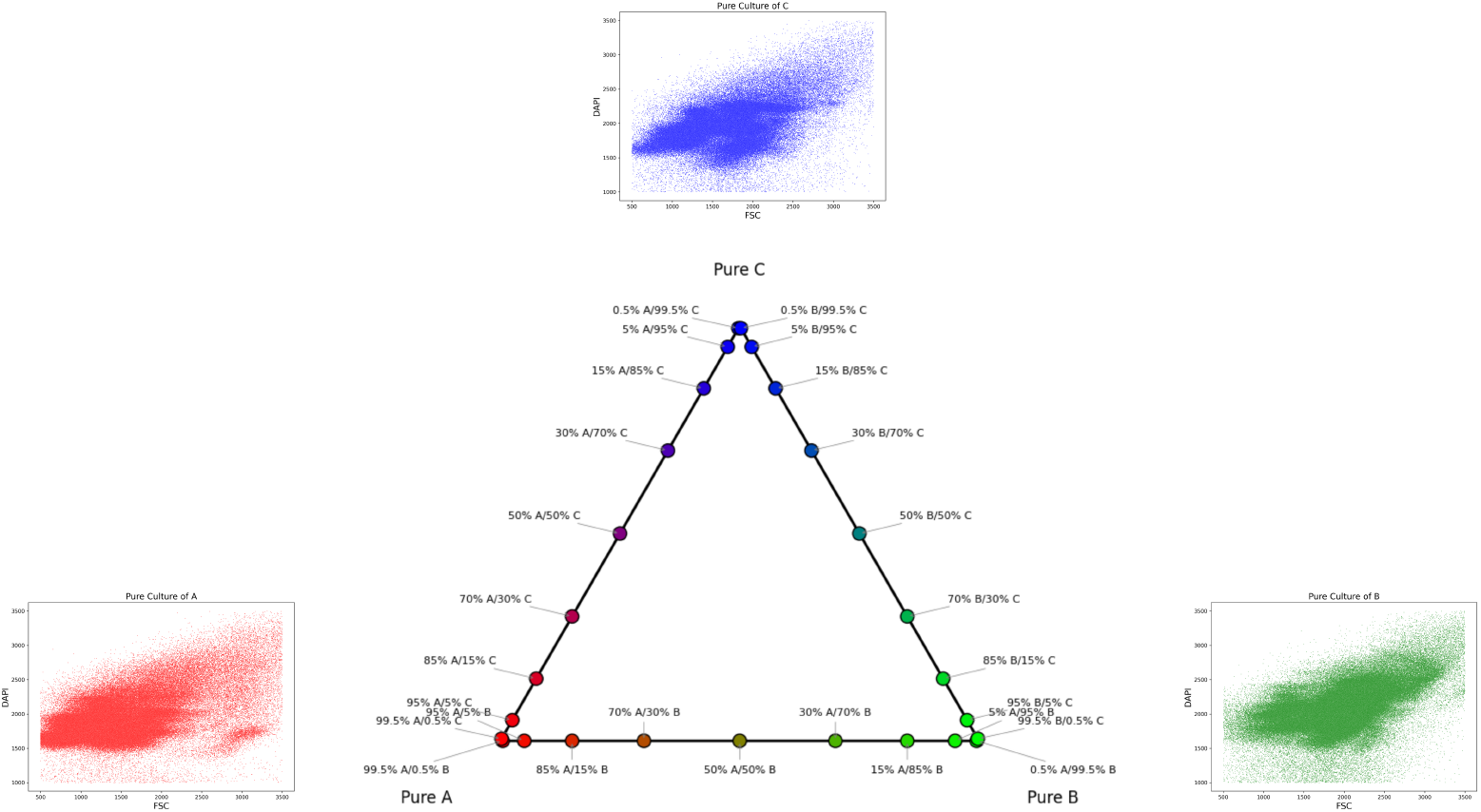
Panel A: Schematic representation of the CHIC benchmark dataset. The dataset comprises three pure bacterial cultures (A, B, and C) and pairwise mixtures at ten distinct ratios (100%, 99.5%, 95%, 85%, 70%, 50%, 30%, 15%, 5%, and 0.5%), each replicated three times, resulting in 90 datasets. The triangular diagram visually represents the varying mixture compositions used. Panel B: Representative flow cytometry scatter plots of pure cultures A (red), B (green), and C (blue), illustrating distinct phenotypic distributions based on Forward Scatter (FSC) and DAPI staining intensity. These pure culture datasets provide the foundation for the benchmarking of computational methods.

For quantitative comparison, we implemented multidimensional scaling (MDS) using pre-computed Wasser-stein distances (convergence threshold: 0.001) and measured accuracy by the frequency with which nearest neighbors in the MDS correctly corresponded to biological replicates.

In order to rigorously evaluate the optimal transport (OT) on bacterial flow cytometry data methods, the benchmark dataset of Koch et al.[20] was used. This dataset consists of three pure (unmixed) samples (A, B, C) and several pairwise mixtures. Each mixture was created at ten different ratios (100%, 99.5%, 95%, 85%, 70%, 50%, 30%, 15%, 5%, and 0.5%), each replicated three times, for a total of 90 data sets (Figure 3). After noise and debris removal, each dataset contained an average of approximately 193,656 cells. Given the uniformity of the data sets with respect to their value ranges, it was determined that no additional preprocessing was necessary. The primary objectives of this benchmarking experiment were to reliably identify replicates and accurately discriminate subtle differences in mixture ratios. To quantitatively compare the methods, we implemented multidimensional scaling (MDS) using pre-computed Wasserstein distances, ensuring convergence with a strict threshold of 0.001. Accuracy was measured by the frequency with which nearest neighbors in the MDS correctly corresponded to biological replicates relative to the total number of comparisons.

We evaluated several distinct computational approaches to benchmark optimal transport methods. As a baseline, we used the original CHIC pipeline as described by Koch et al. [20]. We then explored a local Gaussian Mixture Model (GMM) approach where independent GMMs were fitted to each dataset separately, testing different numbers of mixture components ranging from 5 to 100 (5, 10, 15, 20, 30, 40, 50, 70, 100). Additionally, we implemented a global GMM strategy where a single GMM was fitted to the combined datasets with component counts from 5 to 40 (5, 10, 15, 20, 30, 40), followed by hard clustering and empirical estimation of means and covariances for each dataset. To assess the performance of classical Optimal Transport (OT), we computed Earth Mover’s Distance (EMD) at the individual cell level, employing random subsampling techniques (2,000, 5,000, and 10,000 cells) to address computational complexity. We also evaluated Sinkhorn-regularized OT with entropic regularization (parameter: 0.1) after min-max scaling, using the same subsampling strategy as with classical OT. Finally, we examined a global-to-local GMM approach that combined the strengths of previous methods, using global GMM initialization followed by local refinement with component counts ranging from 5 to 40 (5, 10, 15, 20, 30, 40).

The local GMM approach consistently performed poorest across all component numbers, with accuracy often below the CHIC pipeline baseline (0.5). This under-performance likely resulted from inconsistent component alignment across datasets due to random initialization. The CHIC pipeline demonstrated moderate accuracy (0.5) with adequate performance for balanced mixtures (30%-70%), but struggled with extreme mixture ratios (near 100% or 0%).

Clasical Optimal Transport with EMD showed improved performance reduced datasets. Using just a sample size of 5,000 cells (approximately 2.5% of the original data) exceeded the baseline accuracy, while 10,000 cells achieved approximately 0.60 accuracy. However, the computational costs of this method is extremely high, around 40 times higher than GMM-based approaches.

The global GMM approach consistently outperformes the baseline across all numbers of components, with highest performance (accuracy = 0.66) using 10 mixture components. Further improvement was achieved by global-to-local GMM strategy with 15 components, reaching an accuracy of 0.69. This approach was particularly effective at discriminating extreme mixture ratios (Figure 4) Based on this benchmarking, we selected the global-to-local GMM-OT approach for subsequent analyses.

**Figure 4:**
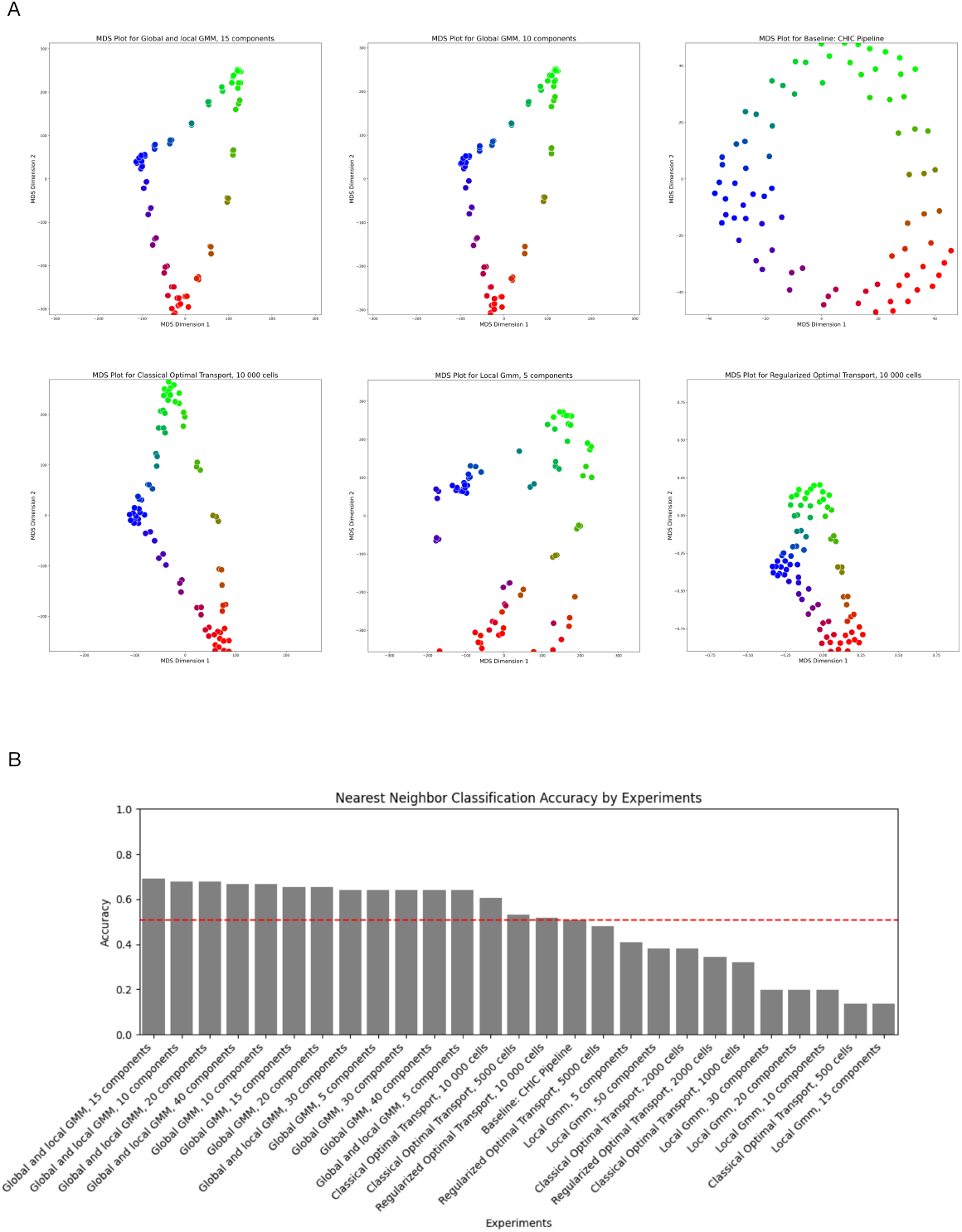
Panel A: Multidimensional Scaling (MDS) plots visualizing Wasserstein distances computed using the best-performing configuration of each OT-based methodology evaluated in this study. From left to right and top to bottom: (i) global-to-local GMM (15 components), (ii) global GMM (10 components), (iii) baseline CHIC pipeline, (iv) classical Optimal Transport with Earth Mover’s Distance using 10,000 cells, (v) local GMM (5 components), and (vi) Sinkhorn-regularized Optimal Transport (10,000 cells). Each point represents an individual dataset, with colors denoting different mixture ratios. Proximity of replicate datasets within each plot reflects the method’s ability to correctly cluster similar samples. Panel B: Summary of nearest neighbor classification accuracy across methods. The global-to-local GMM strategy achieves the highest accuracy, surpassing classical Optimal Transport with random subsampling, local GMM approaches, the CHIC baseline, and Sinkhorn-regularized Optimal Transport. This indicates the enhanced performance of global-to-local GMM initialization in effectively discriminating even subtle differences among complex mixtures while taking computation time into consideration.

### Tracking Bacterial Subpopulation Dynamics over Time

Building on our benchmarking results, which demonstrated thta the global-to-local Gaussian Mixture Model (GMM) approach provides superior accuracy and computational efficiency, we applied this method to track bacterial cell populations over time. We analyzed flow cytometry data from *B. subtilis* cultures collected at six sequential timepoints characterized by optical density (OD). Samples were collected at OD 0.05 (initial), OD 0.5 (at 4.5 hours), OD 1.0 (at 5 hours), followed by a slight decrease to OD 0.92 (at 6 hours), then OD 1.89 (at 8 hours), and finally OD 6.67 (at 24 hours). Manual Gating was performed only performed at OD level 1. These expert-defined gates provided ground truth for temporal comparisons. Consistent preprocessing across all timepoints included noise and debris removal, resulting in 10 well-defined gates: four corresponding to spore subpopulations, one representing transitional cell states, and the remaining gates reflecting distinct vegetative cellular states primarily characterized by chromosome counts and aggregates. We applied logarithmic transformation to all datasets to normalize signal distributions.

We implemented our global-to-local initialization strategy by first fitting a global GMM with 15 Gaussian components to the pooled data across all OD measurements. This global model yielded stable parameter estimates that served as initial values for the local GMMs fitted separately to each timepoint. The choice of 15 components was guided by both prior biological knowledge of *B*.*subtilis* differentiation states and by the empirical performance observed in our benchmarking experiments.

For dynamic tracking, we established the manually-gated OD 1.0 dataset as our reference point and computed pairwise Wasserstein distances to neighboring timepoints. Specifically, forward mappings traced the evolution from OD 1 to 0.92 to 1.89 to 6.67, while backward mappings covered OD 1 to 0.5 to 0.05. From these mappings, we derived optimal transport maps that allowed precise tracking of specific subpopulations, particularly focusing on spore transitions(Figure 5).

**Figure 5:**
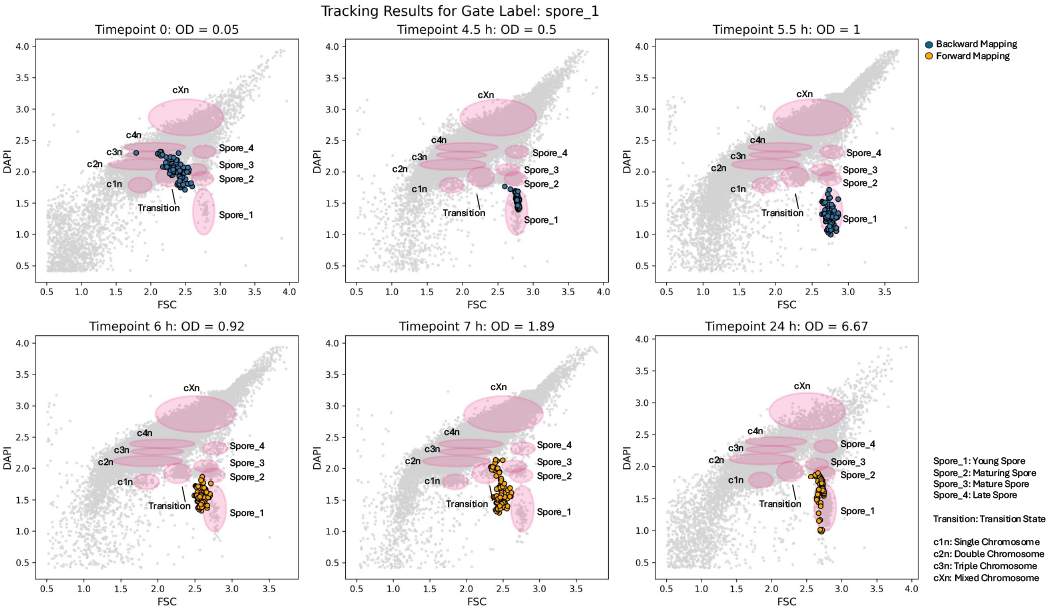
Phenotypic transitions within *Bacillus subtilis* spore populations were monitored from early latent phase (OD = 0.05) through late stationary phase (OD = 6.67). Distinct cell states (spores and vegetative cells differentiated by chromosome count) are visualized as ellipses derived from manual gating. Highlighted here is the temporal evolution of young spores (manual gate: spore 1 from OD 1), tracked incrementally through OT mappings. Blue points represent spores mapped backward in time (towards earlier ODs), while yellow points illustrate forward mappings (towards later ODs). This mapping illustrates the progressive phenotypic transition of spores from an initial inactive and dispersed state into increasingly differentiated subpopulations entering active growth phases.

Our analysis revealed distinct temporal dynamics in spore populations. At the earliest measurement (OD = 0.05), spores exhibited high phenotypic heterogeneity, appearing dispersed across multiple gates. This dispersion indicates anunstable state characteristic of either inactive spores or early-stage germination heterogeneity. As cultures progressed through lag phase, these initially dispersed spores consolidated into more stable, well-defined subpopulations with reduced phenotypic variance. This finding aligns with established descriptions of bacterial spores transitioning from dormancy to environmental sensing and germination initiation phases.

Between OD values of 1.0 and 0.92 (time points 5.0 and 5.5 hours), we observed minimal phenotypic shifts, indicating a stable cellular state over this short interval. However, during the subsequent transition from OD 0.92 to OD 1.89, we detected a substantial expansion and diversification of spore populations into clearly differentiated subpopulations.

At the final time point (OD = 6.67), newly emerging spore subpopulations suggested further cellular differentiation as the culture entered stationary phase. Our incremental OT-based tracking enabled detection of subtle phenotypic shifts that progressively accumulated into significant diversification at higher optical densities. Kathrin said this.

### Integration of B. subtilis flow cytometry and scRNA-seq data

Beyond temporal tracking, we applied our Optimal Transport (OT) framework to integrate unpaired single-cell RNA sequencing (scRNA-seq) and flow cytometry measurements from *B*.*subtilis* cultures grown under similar conditions. This integration enables the imputation of gene expression profiles onto flow cytometry data, creating a unified view of bacterial states across both phenotypic and transcriptomic dimensions. We preprocessed scRNA-seq datasets using the `BacSc` pipeline [25], which is specifically designed to address bacterial transcriptomic challenges including extreme sparsity and high technical noise. After quality control and preprocessing, our scRNA-seq dataset contained 2,784 cells. To facilitate meaningful integration with flow cytometry, we performed dimensionality reduction using Principal Component Analysis on the scRNA-seq data, retaining the top three principal components. This three-dimensional representation balanced dimensionality reduction with information preservation, while conveniently matching the three available flow cytometry parameters: Forward Scatter (FSC), Side Scatter (SSC), and DAPI staining intensity. We log-transformed the flow cytometry data to ensure compatible numerical scales between modalities, after first removing mock communities and noise.

We then fitted Gaussian Mixture Models (GMMs) with 15 mixture components separately to the scRNA-seq and flow cytometry datasets (Figure 6 A). The number of components was chosen to match our temporal analysis for consistency. Pairwise Wasserstein distances were computed between these GMMs to facilitate alignment using Optimal Transport, creating a common embedding space that positioned flow cytometry cells according to the distribution of scRNA-seq-derived gene expression patterns (Figure 6 B). The OT algorithm converged after approximately 100 iterations, indicating robust alignment between the modalities. Within this aligned embedding, we imputed gene expression for each flow cytometry cell by identifying its nearest scRNA-seq neighbor, effectively transferring detailed transcriptomic profiles to phenotypic measurements.

**Figure 6:**
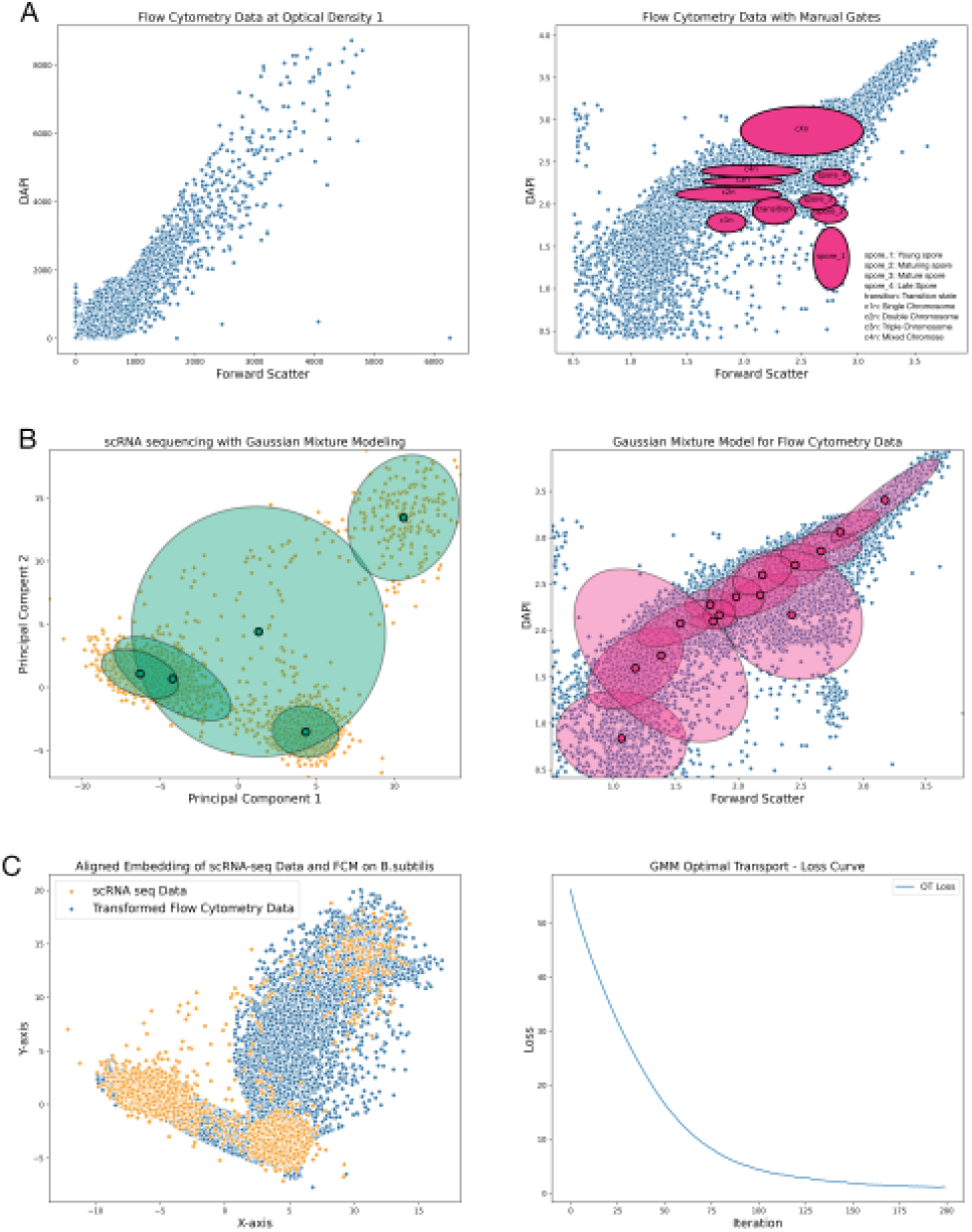
Panel A: Flow cytometry data collected at Optical Density (OD) = 1.0. Left: Raw data distribution before log-transformation, following removal of debris and mock communities. Right: Data after log-transformation and manual gating, with identified bacterial subpopulations annotated. Ellipses indicate manually defined gates representing spores at various stages (young to mature) and cellular states characterized by chromosome counts. Panel B: Gaussian Mixture Model (GMM) fits applied independently to scRNA-seq (left) and flow cytometry (right) datasets. The left plot shows GMM fit on scRNA-seq data after dimensionality reduction while the right plot depicts the corresponding GMM fit on transformed flow cytometry data (Forward Scatter vs. DAPI). Panel C: Left: Aligned embedding resulting from Gaussian Mixture Model-based Optimal Transport (GMM-OT), integrating scRNA-seq (orange points) and transformed flow cytometry data (blue points). Right: Convergence of the OT algorithm shown through loss reduction, stabilizing after approximately 100 iterations.

To validate the biological relevance of our gene expression imputation, we examined four well-characterized B. subtilis marker genes with distinct functional roles (Figure 7). The manual gating framework established in Figure 6A provided a reference by dividing cells into vegetative (c1n to c4n), aggregated (Cxn), and spore-forming (spore1 to spore4) populations, along with transitional states.

**Figure 7:**
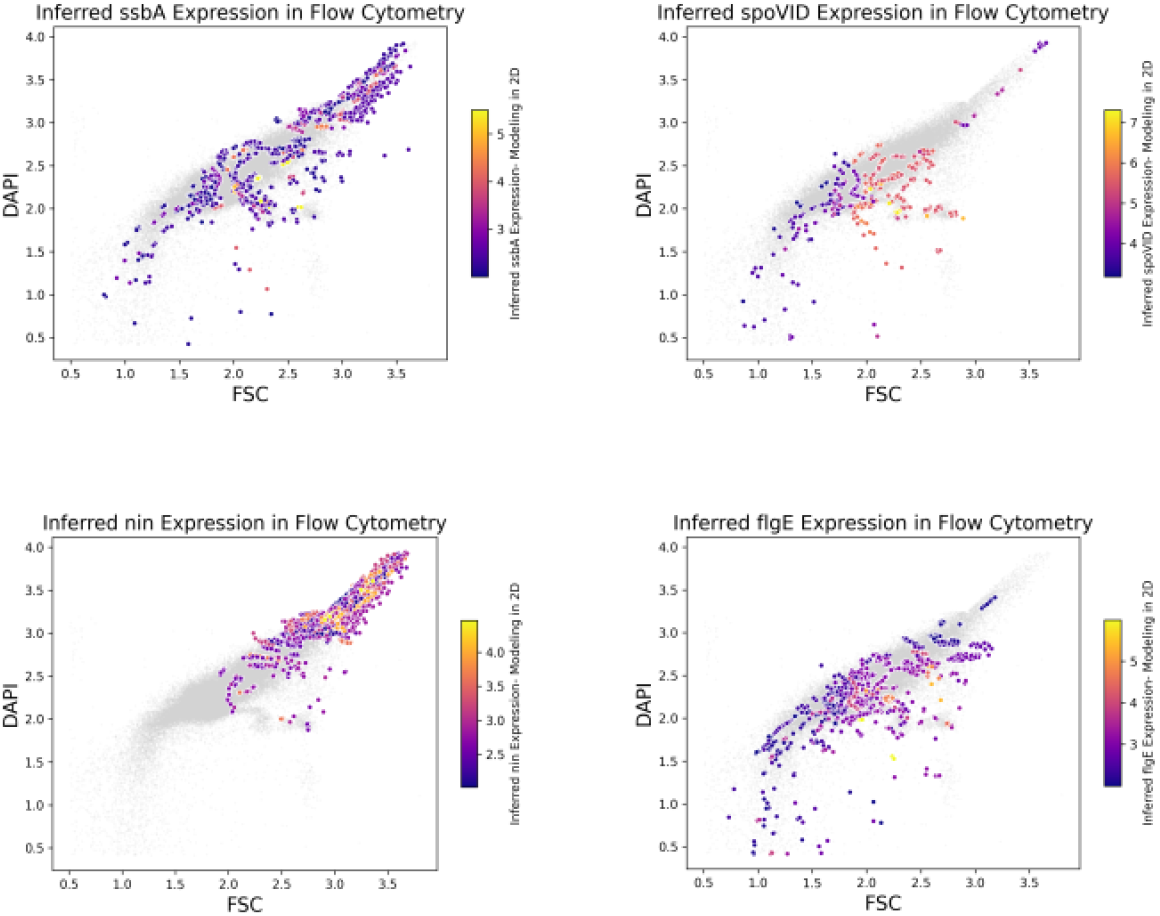
Comparison of inferred gene expression for four characteristic *B. subtilis* marker genes. Expression levels are shown as a color gradient overlaid on flow cytometry measurements of FSC and DAPI. Top left: *ssbA* (vegetative cell marker) shows widespread expression with higher intensity in vegetative and aggregated regions. Top right: *spoVID* (sporulation marker) exhibits highest expression in spore-associated gates. Bottom left: *nin* (DNA regulation) localizes to high FSC/DAPI subpopulations associated with aggregated cells. Bottom right: *flgE* (motility) shows scattered expression predominantly in vegetative cells. Manual gates are overlaid for reference.

The single-stranded DNA binding protein gene *ssbA* (Figure 7, top left) plays a critical role in DNA replication, repair, and recombination, essential for maintaining genomic stability [26, 27]. As expected, imputed *ssbA* expression appeared widespread throughout the flow cytometry plot, with highest expression in vegetative cell gates (c1n-c4n) and aggregated cells, consistent with its role in actively replicating cells.

The sporulation-associated gene spoVID (Figure 7, top right) encodes a protein that anchors spore coat proteins during spore maturation [28, 29]. Our imputation showed spoVID expression predominantly within spore-associated gates characterized by intermediate to low DAPI intensities and intermediate to high FSC values. Intermediate expression levels in transition states accurately reflected the expected expression pattern in cells progressing toward sporulation.

The DNA regulation gene nin (Figure 7, bottom left), which encodes an inhibitor of NucA activity involved in DNA degradation during competence induction [30], showed localized expression in subpopulations with higher FSC and DAPI intensities. This pattern corresponds to cells with increased DNA content or replication activity (cXn), consistent with nin’s role in preventing DNA degradation during competence development.

Finally, the flagellar filament gene flgE (Figure 7, bottom right) [31] showed expression primarily in vegetative cells corresponding to manual gates c1n through c4n. The scattered expression pattern throughout these gates aligns with the heterogeneous nature of motility gene expression in bacterial populations.

The gene expression patterns revealed by our imputation approach show remarkable concordance with known biological functions and cellular states in B. subtilis. This validation demonstrates that our GMM-OT framework successfully bridges flow cytometry and scRNA-seq modalities, enabling the integration of rich transcriptomic information with high-throughput phenotypic measurements.

### Three-dimensional integration of B. subtilis flow cytometry and scRNA-seq data

Bacterial flow cytometry data are inherently limited by the number of channels measured. In our dataset, three channels are available, namely forward scatter (FSC), side scatter (SSC) and DAPI. To improve the biological resolution, we integrated these measured dimensions with the latent dimensions. Although the pipeline suggests four optimal latent dimensions, we were limited to incorporating only the top three due to the inherent dimensional limitations of our cytometry measurements.

While manual gates are not available in this case and therefore interpretation is limited, Figure 8 demonstrates how our approach can be used for these cases. The top row shows the inferred expression of *spoIVA*. The left panel shows DAPI versus side scatter, while the right panel shows side scatter versus forward scatter. In the upper right panel, high levels of *spoIVA* expression are characterized by higher Forward Scatter values relative to Side Scatter values.

**Figure 8:**
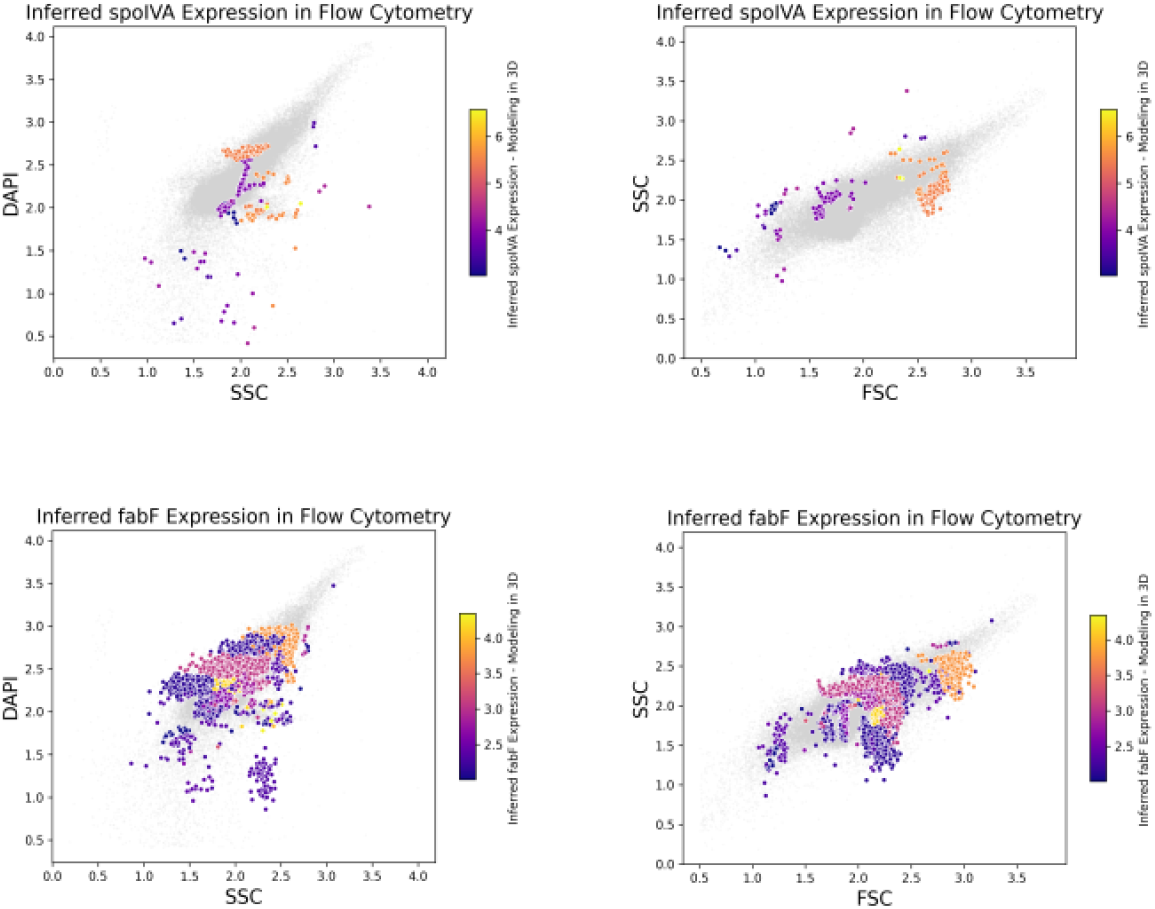
Inferred gene expression patterns for sporulation and vegetative cell markers visualized across different cytometry parameters. Top row: *spoIVA* expression in DAPI vs. SSC (left) and SSC vs. FSC (right) plots. The sporulation-associated *spoIVA* gene shows highest expression (yellow) in cells with intermediate DAPI intensity, higher FSC and medium SSC values. Bottom row: *fabF* expression in DAPI vs. SSC (left) and SSC vs. FSC (right) plots. The metabolic gene *fabF* shows a sectional distribution pattern typical of vegetative cells, with highest expression concentrated at FSC values around 2.2 and SSC values around 2.0.

*spoIVA* encodes a protein that is critical for sporulation, specifically for spore coat formation. It is one of the key proteins required for proper assembly of the spore coat and is essential for the sporulation process in *Bacillus subtilis*. [32]. During sporulation, *spoIVA* functions in the mother cell compartment and is responsible for the inward folding of spore coat proteins, facilitating the encapsulation of the developing forespore. While these patterns are consistent with the known biology of *spoIVA*, further conclusions are limited by the lack of manual gates in three dimensions. The highest *spoIVA* expression can be found in cells with high FSC but medium SSC values, as well as in regions with intermediate DAPI intensities. Although there are limitations, we observe typical features associated with spore cells.

The lower panel of Figure 8 shows the inferred gene expression of *fabF*. This gene encodes *β*-ketoacyl-ACP synthase II, an enzyme involved in the elongation step of fatty acid synthesis. The enzyme catalyzes a key reaction in the elongation cycle, specifically the addition of two-carbon units to a growing fatty acid chain and textitfabF is predominantly expressed in vegetative cells [33]. Examining the SSC vs. DAPI plot reveals a distinctive sectional pattern of *fabF* expression. Cells with DAPI values between 2.5 and 2.8 exhibit moderate to high expression levels, transitioning to regions of reduced expression, before intensifying again at DAPI values around 3.0. The SSC vs. FSC plot displays comparable distribution patterns, with peak *fabF* expression occurring in cell populations characterized by FSC measurements of approximately 2.2 and SSC measurements of approximately 2.0.

## Discussion

In this study, we systematically evaluated several Optimal Transport (OT)-based computational strategies for alignment and integration of heterogeneous single-cell datasets. Our benchmarking revealed significant performance differences between classical OT (Earth Mover’s Distance), entropically regularized OT (Sinkhorn), and Gaussian Mixture Model-based OT (GMM-OT) for bacterial flow cytometry data. Our results demonstrate that `biscot`, employing GMM-OT combined with carefully designed Gaussian Mixture Model initializations, offer significant computational advantages for handling complex single-cell data.

Benchmarking against the established CHIC datasets [20] showed that classical OT (EMD) provides robust accuracy in distinguishing replicate samples and subtle mixture ratios, even under heavy subsampling. However, the computational burden associated with classical OT limits its applicability in practice, especially for large datasets. Entropic regularization via the Sinkhorn algorithm significantly reduces computational complexity while maintaining acceptable accuracy. However, this approach introduces additional hyperparameter tuning, which may complicate its practical use.

Critically, our results showed that independently fitted local GMMs performed poorly due to inherent instability caused by random initialization and inconsistent component labeling across datasets. To overcome these problems, we introduced and validated a global-to-local GMM initialization approach. By first fitting a global GMM to pooled datasets and then refining local models based on these global parameters, we achieved consistent component alignment and significantly improved accuracy, particularly in biologically relevant scenarios involving unbalanced mixtures. The global-to-local GMM strategy employed in `biscot` emerged as the optimal solution, consistently outperforming classical OT, Sinkhorn OT, and baseline methods.

We applied `biscot` to track temporal dynamics in *Bacillus subtilis* populations. The method effectively captured subtle but biologically significant transitions in spore subpopulations as they transitioned from inactive to active growth. The detailed temporal trajectory constructed by incremental OT mappings provided biologically interpretable insights into cellular differentiation stages, in close agreement with known biological processes governing spore germination cite. This highlights the ability of GMM-OT to robustly quantify microbial population dynamics in longitudinal studies.

The integration of flow cytometry with single-cell RNA sequencing (scRNA-seq) further demonstrated the versatility and biological relevance of `biscot`. By imputing scRNA-seq-derived gene expression profiles to phenotypically defined flow cytometry cells, we enriched traditional cytometric analyses with transcriptomic insights. Validation with known marker genes, including *spoVID, ssbA, nin* and *flgG*, confirmed the biological plausibility of the imputed expression profiles, underscoring the reliability.

Despite these advances, several important limitations remain. The selection of 15 Gaussian components was guided primarily by biological intuition and initial exploratory analyses; rigorous, data-driven optimization of the number of components using systematic criteria, such as cross-validation or information-theoretic approaches, would further enhance model accuracy and interpretability.In the future, additional validations using different biological systems and paired modalities would increase confidence in the generalizability of our OT-based framework.

In summary, our work demonstrates that `biscot` provides a robust, biologically interpretable, and computationally efficient solution for integrating and analyzing complex microbial single-cell data. By bridging phenotypic and transcriptomic modalities, `biscot` has significant potential to advance our understanding of dynamic cellular processes in microbial populations, opening new avenues for detailed characterization of cellular states, transitions, and functional adaptations.

## Computational Environment and Reproducibility

All analyses were performed using Python 3.12.9. The GMM estimation utilized `sklearn.mixture` (v1.6.1)[23], while Optimal Transport calculations were implemented using the `POT` [34] library (v0.9.5). Complete source code, including configurations and preprocessing pipelines, is available at [https://github.com/bio-datascience/flow-alignment] for full reproducibility.

## Data Availability

## Acknowledgements

We thank Adam Z. Rosenthal (UNC) for in-depth advice and help with the B. subtilis scRNA-seq data. C.L.M. acknowledges core funding from the Institute of Computational Biology, Helmholtz Zentrum München. This work was supported by the Deutsche Forschungsgemeinschaft (DFG) within the SPP 2389 priority program “Emergent Functions of Bacterial Multicellularity” (Grant number: 503905203) awarded to S.M and C.L.M.

## Competing Interests

The authors declare no competing interests.

## Notes

### Competing Interest Statement

The authors have declared no competing interest.

